# G protein subtype preference dictates paroxetine-enhanced serotonin receptor response in normal breast epithelial cells

**DOI:** 10.1101/2025.06.10.658790

**Authors:** Máté Lengyel, Péter Árkosy, Iván P. Uray

## Abstract

In the healthy mammary gland serotonin (5-HT) has a key role in shaping tissue morphology and regulating lactation and involution. Serotonin re-uptake inhibitors (SSRIs) alter the distribution of 5-HT across the cell membrane, but how increased extracellular 5-HT may impact breast cell homeostasis is unclear. We demonstrate that paroxetine reduces 5-HT levels in immortalized primary breast epithelial (HME-hTert) cells, mitigates free oxygen radical formation and decreases cell migration and proliferation. Accordingly, pathways related to cell cycle and DNA damage repair were underrepresented in the transciptomic profile of paroxetine-treated cells. On the other hand, enriched transcipts overrepresented genes affecting neural transmission and GPCR signaling, suggesting an increase in 5-HT receptor (5HTR) activation. As 5-HT induced the levels of both cAMP and inositol triphosphate (IP3), the contribution of individual 5HTRs were decyphered using receptor-selective agonists and antagonists. 5HTRs coupled to each Gα protein subtype were expressed and functional in HME-hTert cells. The activation of G_S_-coupled receptor 5HTR7 and antagonists of G_i_- and G_q_-coupled receptors 5HTR1D and 5HTR2B generally suppressed proliferation. The induction of cAMP levels by 5-HT was reduced by the 5HTR7 antagonist and IP3 induction was blocked by an 5HTR2B inhibitor, while the 5HTR1D antagonist further increased cAMP levels induced by 5-HT. Paroxetine-dependent growth suppression was reversed by inhibitors of G_S_-coupled 5HTR7, protein kinase A or adenylyl cyclase, and agonists of G_i_ or G_q_-coupled receptors. These results suggest that the anti-proliferative responses to 5-HT in non-malignant breast cells align with the G protein preference of the receptors, and reveal potential benefits of repurposing receptor subtype-selective agents and SSRIs for cancer risk reduction.

**Graphical abstract:** 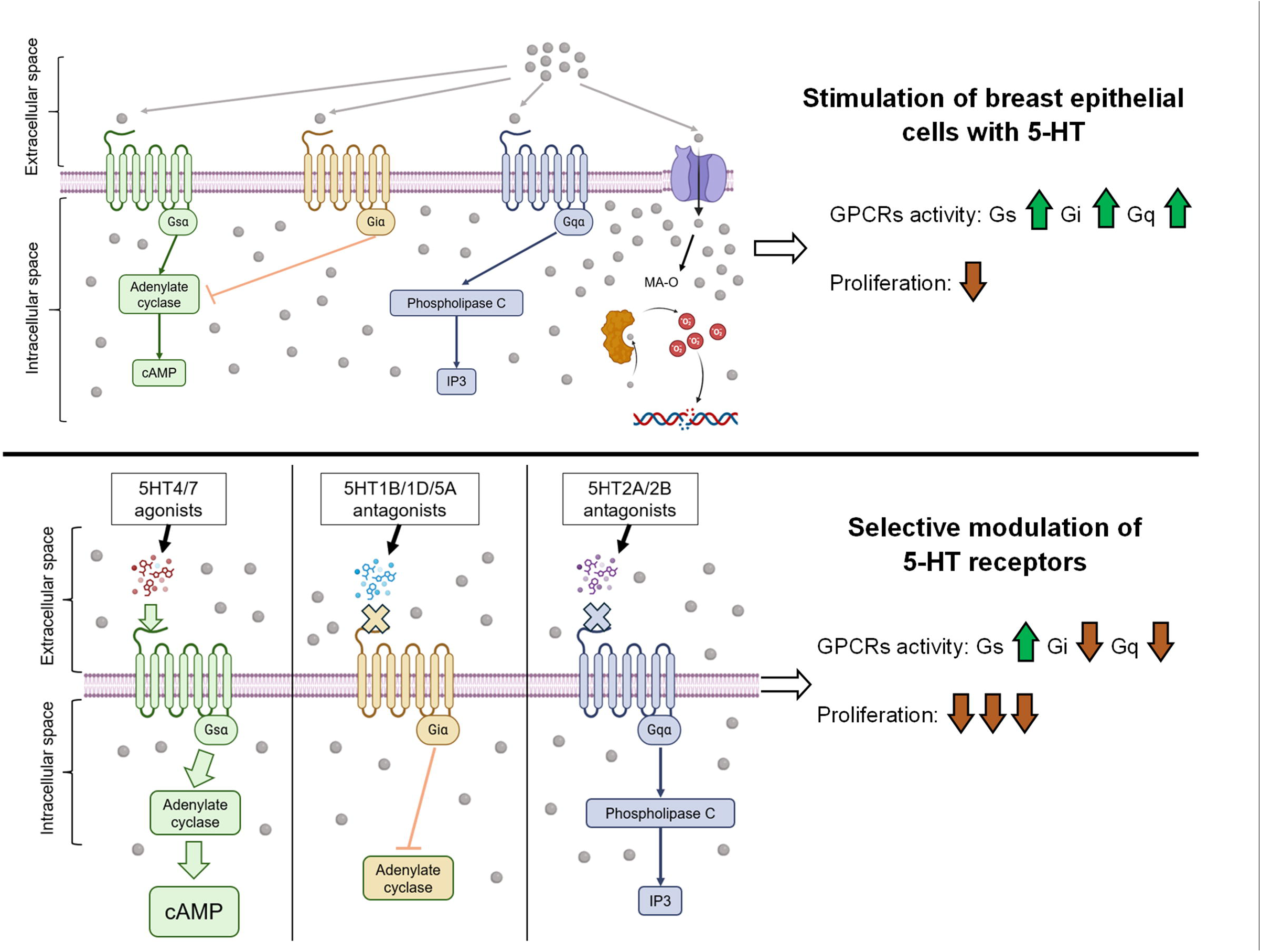

## Introduction

Serotonin (5-HT) as a neural monoamine neurotransmitter regulates mood, memory, circadian cycle and behavior, as well as respiratory and thermoregulatory networks (1, 2). However, mounting evidence indicates that 5-HT produced outside the nervous system has relevant paracrine effects on normal and transformed cells (3-5). In the physiological regulation of the mammary gland serotonin stimulates lactation, initiates involution and allows the mammary gland to drive the mobilization of calcium from the skeleton (6, 7). Disruptions in neuroendocrine homeostasis or a balanced microbiome have marked, but ill-specified impact on oncogenesis and growth (8, 9). In disease, increased serotonin levels were reported in association with advanced breast tumors (10) and 5-HT is often considered a predictive marker for breast cancer (11) (12).

Extracellular serotonin signals are transmitted by 13 metabotropic 5-HT receptors (5HTRs 1-7) and a ligand gated ion channel, 5HTR3 (13). Because all 5HTRs share the same ligand, and specific tissues express distinct levels and combinations of these receptors, defining specific pathways and physiological roles for each of these serotonin receptors has been a challenging task. The signaling events following the activation of a diverse array of serotonin receptors has been studied in normal and cancer cells, and linked with the regulation of cell proliferation and apopotosis (3, 14, 15). Conversely, serotonin response modulators are expected to arise as novel or repurposed therapeutic options acting through 5HTRs (16-18).

In addition to the ability to synthesize 5-HT by the TPH1 enzyme, breast epithelial cells can also collect serotonin from the extracellular space using the sodium:serotonin symporter SERT (SLC6A4) (19). In the synaptic space the inhibition of 5-HT uptake by serotonin re-uptake inhibitors (SSRIs) serves the potentiation and extension of the postsynaptic serotonin signaling. However, in epithelial cells the influence of SSRIs on 5HTR-dependent responses have not been elucidated. We hypothesized that, SSRIs might enhance receptor-dependent signaling in breast epithelial cells similar to neurons, and the type of response will be dictated by the affected receptors and second messengers.

In this study we examined the impact of the SSRI paroxetine on 5-HT distribution and signaling. We identified the serotonin receptors involved and the associated G proteins modulating downstreamm signaling, cell proliferation and motility in response to excess 5-HT in the extracellular space. A library of receptor subtype-specific agonists and antagonists delineated growth inhibitory pathways utilizing 5-HT receptors. We show that the activation of G_sα_ and antagonists acting through the G_i_ and G_q_ are inhibitory to cell proliferation, while G_q_ and G_i_ agonists have little impact. Our data suggest that G protein subtype preference is the key determinant in breast cell response to 5-HT.

## Results

### 5-HT and paroxetine inhibit the proliferation of immortalized primary breast epithelial cells

To evaluate the effects of serotonergic stimulation on breast epithelial cell growth, 2D proliferation assays were performed using HME-hTert cells treated with the selective 5-HT reuptake inhibitors (SSRIs) citalopram, sertraline, and paroxetine. Among these, paroxetine, the most potent SSRI tested, significantly reduced cell counts in a dose-dependent manner at or above 1µM concentrations (Figure 1A). Consistent with the antiproliferative effects observed with paroxetine, siRNA-mediated knockdown of SERT resulted in a significant decrease in total cell counts (Figure 1B). Furthermore, 5-HT alone inhibited proliferation in a dose-dependent manner, and this effect was significantly enhanced when 5-HT was combined with paroxetine (Figure 1C). To assess the effects of paroxetine and 5-HT on anchorage-independent cell growth, 3D breast epithelial spheroids were treated with 5-HT or paroxetine for one week. Both agents exhibited a marked reduction in spheroid volume, further emphasizing the inhibitory effect of serotonergic stimulation on proliferation (Figure 1D). In addition, in wound healing assays paroxetine significantly reduced cell migration compared to vehicle, indicating its impact on cell adhesion and motility (Figure 1E). As a major factor regulating cell motility and matrisome-remodelling in HME-hTert cells, ZG16B was found to be suppressed by paroxetine, either alone or in combination with bexarotene (Supplementary figure 1) (20).

**Figure 1.**
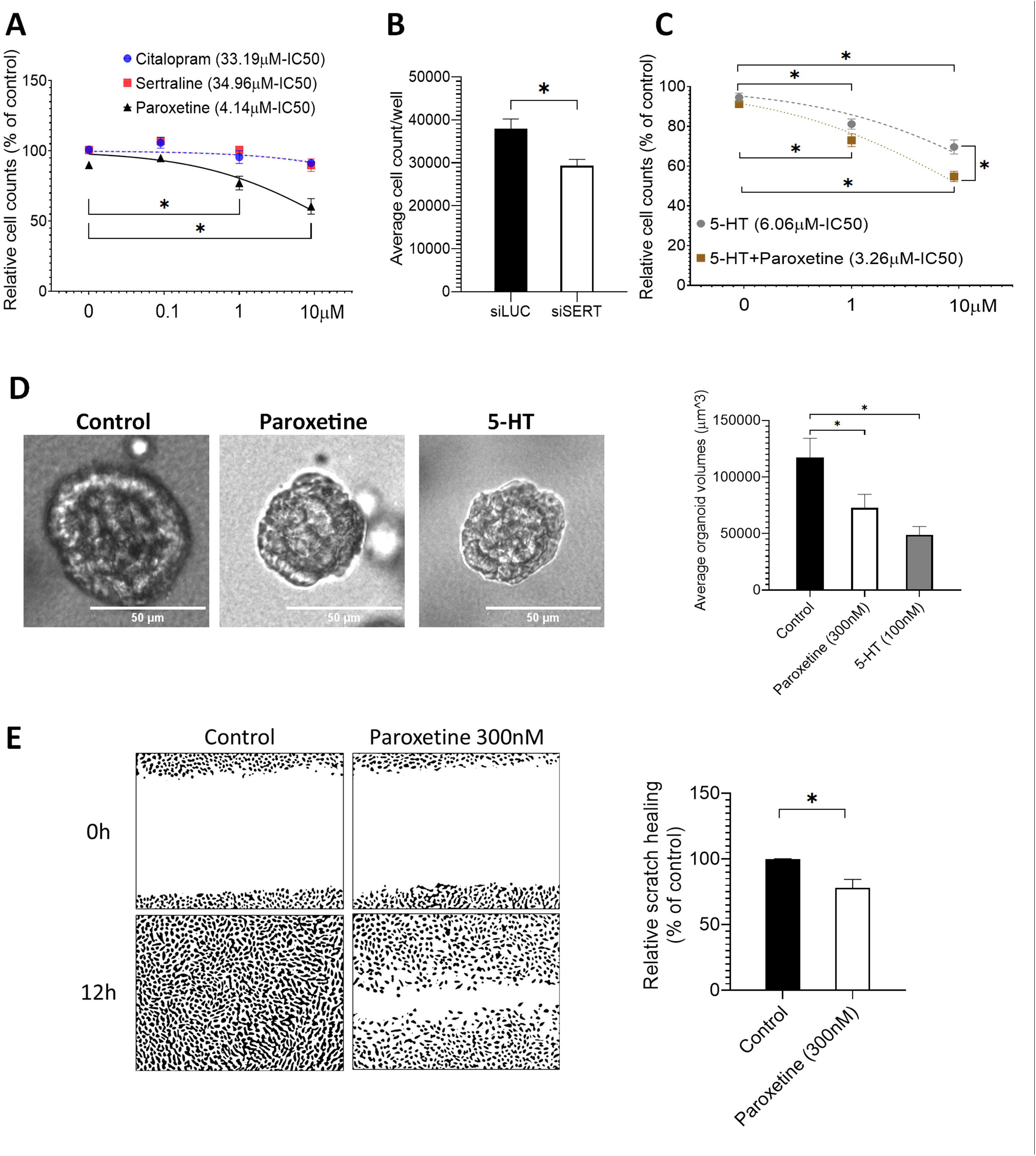
Proliferation of normal breast epithelial cells treated with paroxetine or serotonin (5-HT) assessed in 2D and 3D growth assays. **A**. Dose-response of HME-hTert cell proliferation to SSRIs. 2D growth was assessed upon treatment with SSRIs applied at 0.1, 1 or 10 μM. **B**. Cell proliferation after siRNA-mediated knock-down of SERT. siRNA targeting luciferase was used as negative control. **C**. Comparison of the antiproliferative effect of 5-HT alone or combined with paroxetine on HME-hTert. Cells received control or paroxetine (1μM) pre-treatment followed by 5-HT at 1-10uM concentrations. **D**. Comparison of the antiproliferative effects of 5-HT and paroxetine in HME-hTert-derived epithelial organoids grown in Matrigel. **E**. Assessment of HME-hTert cell migration and adhesion upon paroxetine treatment. Cells were plated on Ibidi 24-well plates with inserts for wound healing assays and cultured until reaching confluency. After removal of inserts vehicle or paroxetine were added and open surface areas were compared at 0 and 12 hours post-treatment.

### 5-HT uptake, ROS production and DNA damage are attenuated by paroxetine

Immunocytochemical labeling of 5-HT revealed detectable intracellular 5-HT levels in untreated HME-hTert cells. In keeping with the activity of functional serotonin re-uptake pumps, paroxetine treatment significantly reduced intracellular 5-HT content (Figure 2A), while SERT levels remained unaffected, as shown by immunocytochemical analysis of SERT expression in HME-hTert cells (Supplementary figure 2). Serotonin is broken down by the activity of the monoamine oxidase A (MAO-A) enzyme, associated with the generation of reactive oxygen species (ROS). Intracellular free radicals, in turn, may cause DNA damage and the formation of γH2AX foci (Figure 2B).

**Figure 2.**
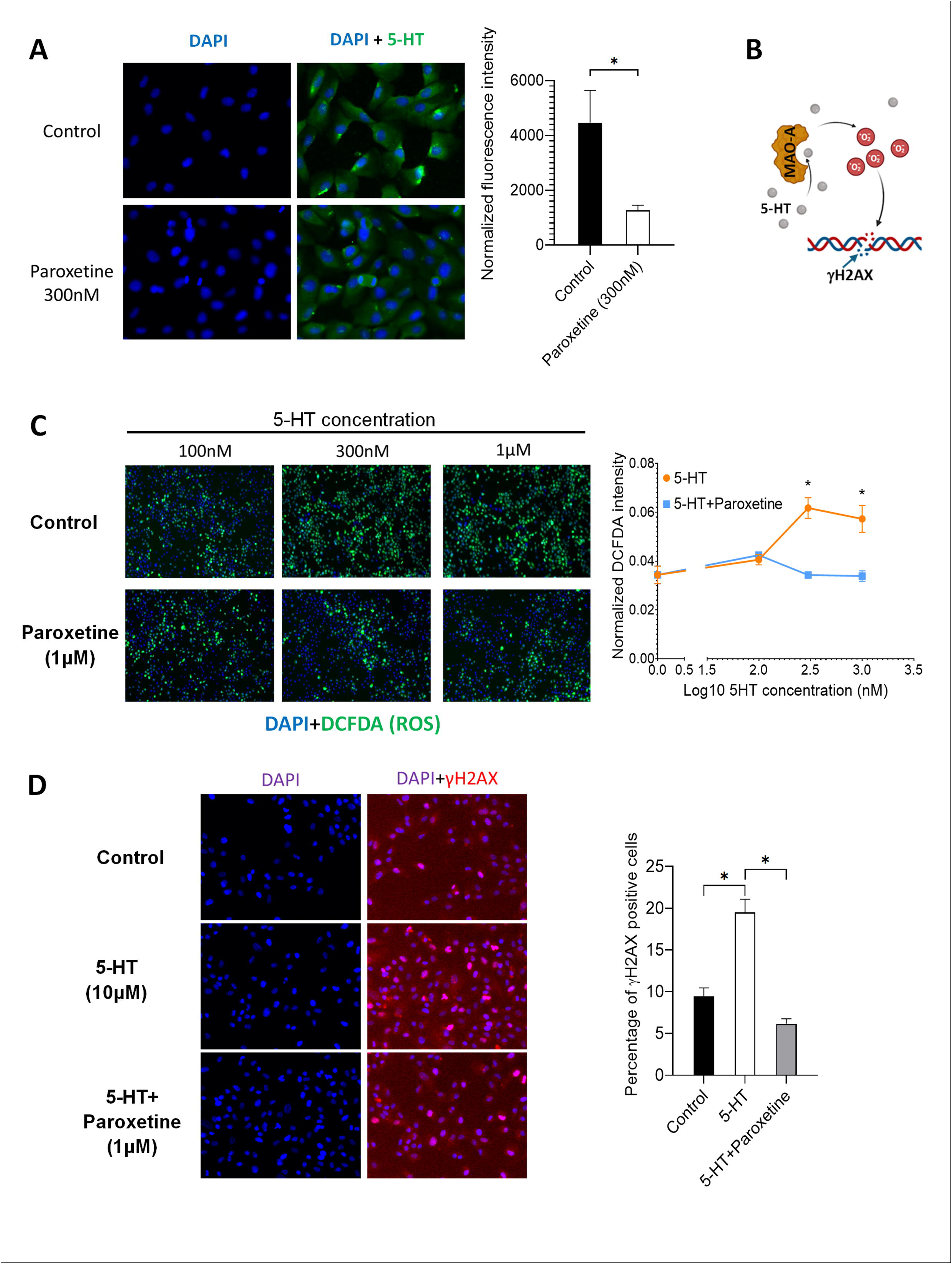
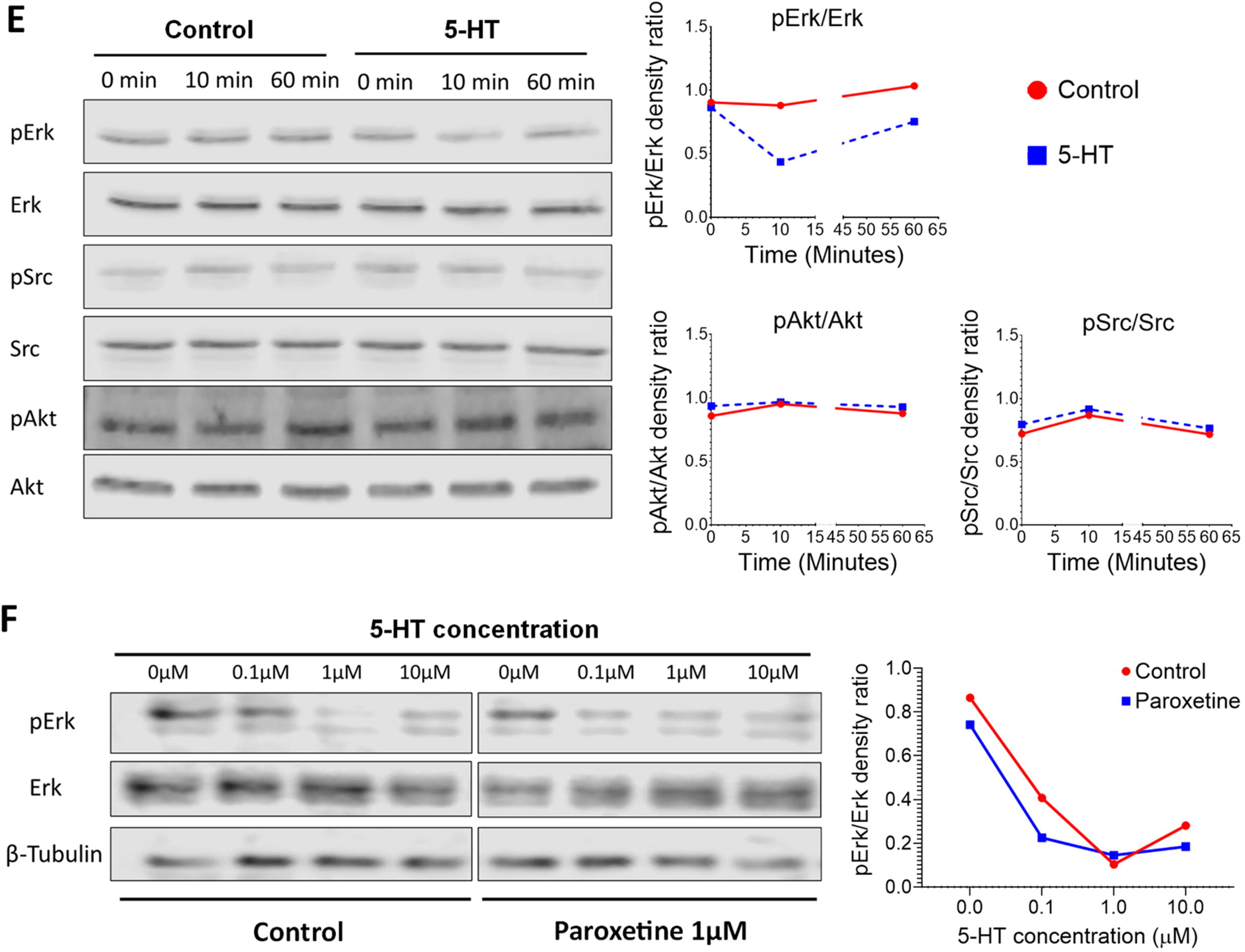
The impact of SERT activity and expression on 5-HT levels, free oxígen radical (ROS) production, DNA damage repair, mitogenic signaling and breast cancer survival. **A**. Quantification of intracellular 5-HT content upon paroxetine treatment (300 nM) in HME-hTert. by immunolabelling (green: 5-HT, blue: DAPI). **B**. Model for the formation of free oxygen radicals as a byproduct of 5-HT breakdown by MAO-A, and the resulting DNA breaks inducing γH2AX foci. **C**. Comparison of ROS production in cells treated with 5-HT alone or combined with 1 μM paroxetine by DCFDA staining. Oxidized DCFDA intensities were normalized on cell counts based on DAPI staining (green: DCFDA, blue: DAPI) **D**. Quantitation of double-stranded DNA breaks after treatment by 5-HT (10 μM) alone or 5-HT+paroxetine (1 μM.) Cell fractions positive for double stranded DNA-breaks were determined based on γH2AX marker (red) positivity, total cell counts were determined based on DAPI staining (blue). **Figure 2E-F**. Western-blot analyses of mitogenic kinase phosphorylation upon 5-HT treatment. Representative images (left panel) illustrate Erk, Src, and Akt phosphorylation 10 or 60 minutes after serotonin treatment (E). Dose-dependence of Erk activation by 5-HT. Representative images (left panel) display Erk phosphorylation in response to increasing 5-HT concentrations after vehicle (control) or paroxetine pre-treatment (F). The line graphs on the right show the ratios of phosphorylated forms normalized to total proteins in the blots shown. The densitometric evaluation was performed in ImageJ.

As expected, 5-HT treatment increased the production of reactive oxygen species (ROS), however, this effect was diminished by paroxetine-mediated inhibition of SERT, as measured by DCFDA fluorescence (Figure 2C). In parallel, elevated γH2AX levels indicated the increased formation of double-stranded DNA breaks upon 5-HT exposure, while paroxetine treatment suppressed this 5-HT-mediated elevation in γH2AX positivity in HME-hTert cultures (Figure 2D).

To elucidate the signaling events associated with the antiproliferative effects of 5-HT and paroxetine on HME-hTert cells, the activation of Erk, Src, and Akt was assessed upon 5-HT treatment. Reduced Erk phosphorylation was detected (average decrease >40%, p<0.05) within 10 minutes of 5-HT treatment, but Src and Akt phosphorylation levels were not affected (Figure 2E). In dose-response experiments 5-HT alone induced Erk dephosphorylation at concentrations as low as 0.1µM (mean suppression 46%, p<0.05, Figure 2F). Moreover, pre-treatment with paroxetine followed by 5-HT exposure further reduced Erk phosphorylation. Clinically, the SLC6A4 gene coding for the SERT protein is amplified in 3-4% of breast cancers (TCGA and METABRIC). Cases with high SERT expression are associated with less favorable prognoses, compared to low expressors (HR=1.32, Supplementary Figure 3).

Paroxetine-induced transcriptomic changes assessed by RNA-sequencing showed a significant underrepresentation of pathways associated with cell cycle regulation and checkpoints, DNA repair and replication, senescence, protein folding and stress response. Gene Set Enrichment Analysis (GSEA) using the Reactome database gene-sets indicated the attenuation of pathways related to G1-S DNA damage checkpoints, pErk-mediated gene expression regulation, p53 stabilization, the regulation of cyclins, APC and the replication origin complex (Figure 3, representative GSEA plots shown in the left panel). Conversely, GSEA gene sets indicated increased activity in neural transmission, increased cell migration and adhesion and a modulation of GPCR signaling (Figure 3). Notably, within these categories, key enriched pathways include the synthesis of IP3 in the cytosol, effects of PIP2 hydrolysis, G-alpha 12/13 signaling events, the Rho GTPase cycle, collagen formation, and integrin cell surface interactions (Representative GSEA plots shown on the left side of Figure 4.)

**Figure 3.**
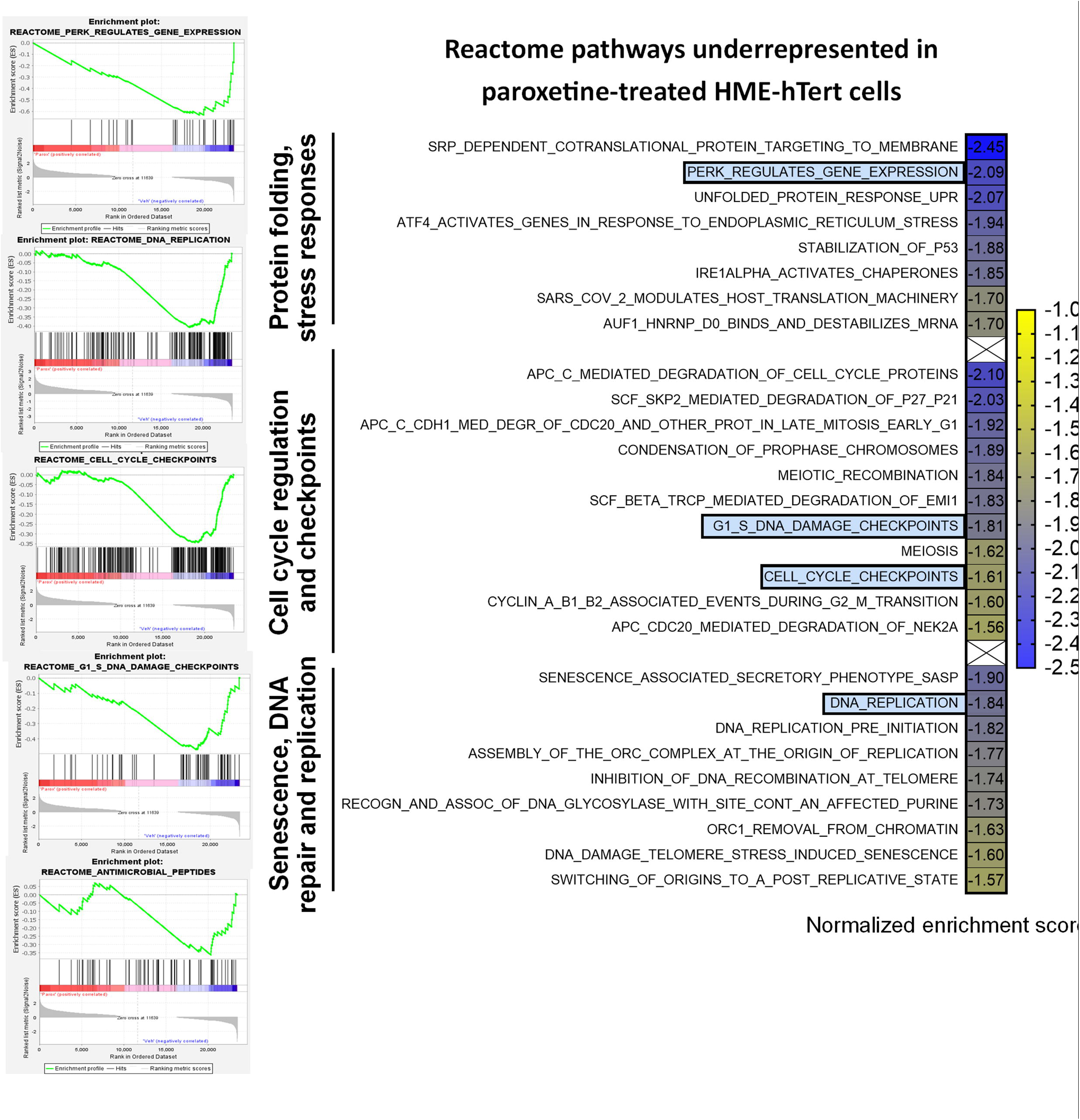
Reactome pathways underrepresented in HME-hTert cells treated by paroxetine over control. GSEA of RNA-seq data obtained from control or paroxetine treated HME-hTert cells was performed using Reactome pathways. Filtering of identified pathways was performed by applying p<0.05 and FDR q-value<0.125 thresholds. The color scale represents normalized enrichment score (green: low enrichment score, red: high enrichment score)

**Figure 4.**
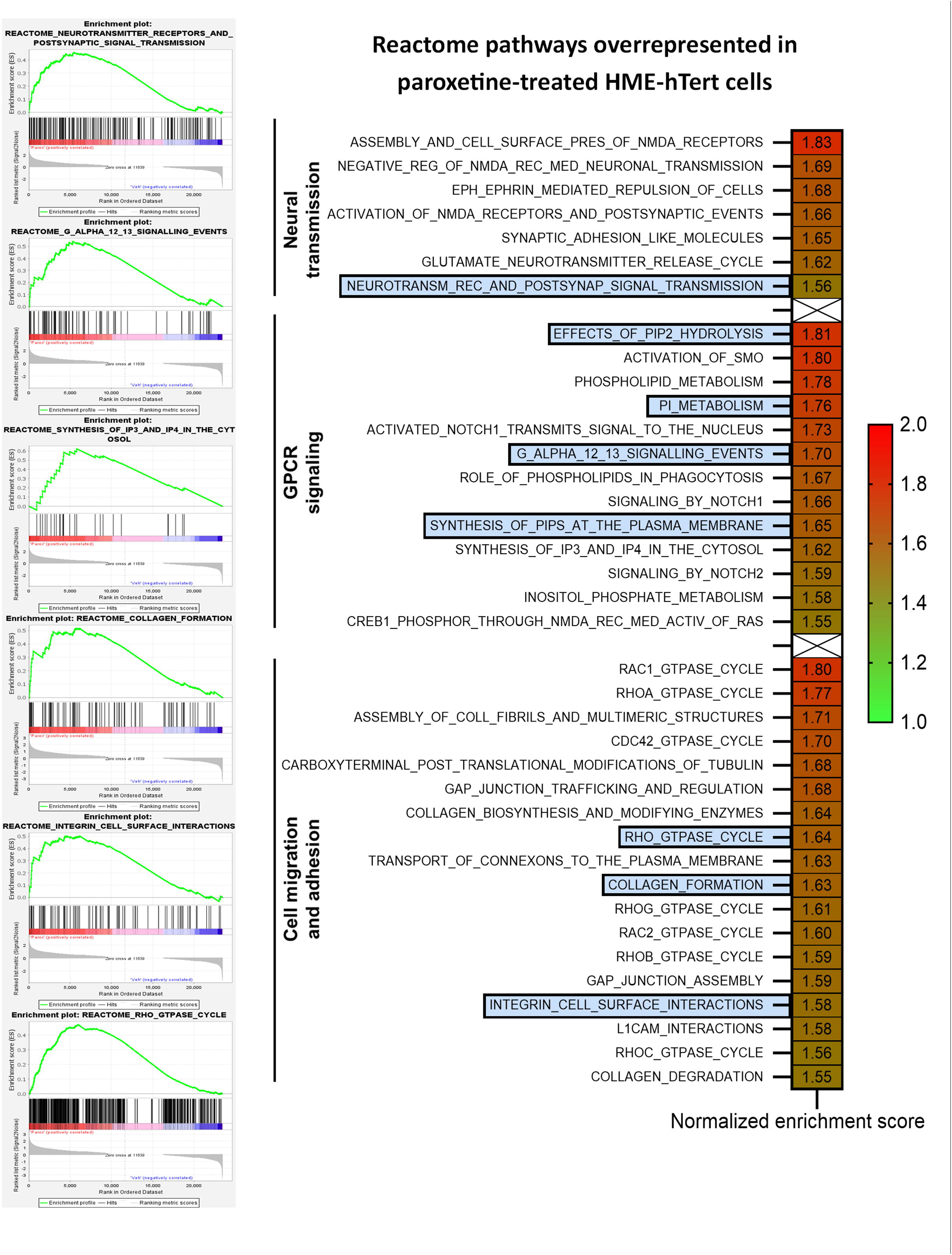
Reactome pathways enriched in HME-hTert cells by paroxetine vs control treatment. Gene set enrichment analysis (GSEA) of RNA-seq data obtained from paroxetine or control treated HME-hTert cells was performed using Reactome pathways, filtering of identified pathways was performed by applying p<0.05 and FDR q-value<0.125 thresholds. The color scale represents normalized enrichment score (green: low enrichment score, red: high enrichment score)

### 5-HT and paroxetine induce 5HTR-specific signaling events in HME-hTert cells

RNA-seq analysis of untreated HME-hTert cells revealed the expression of multiple 5-HT receptor isoforms at varying levels, among which 5HTR7 mRNA was the highest (Figure 5A). RT-qPCR confirmed robust 5HTR7 expression, while 5HTR2B and 5HTR1D exhibited moderate expression levels (Figure 5B). To assess receptor functionality, levels of cAMP and IP3 second messengers were quantified following 5-HT treatment. 5-HT, as the endogenous ligand of 5-HT receptors, was capable of inducing both cAMP and IP3 production (Figure 5C, D).

**Figure 5.**
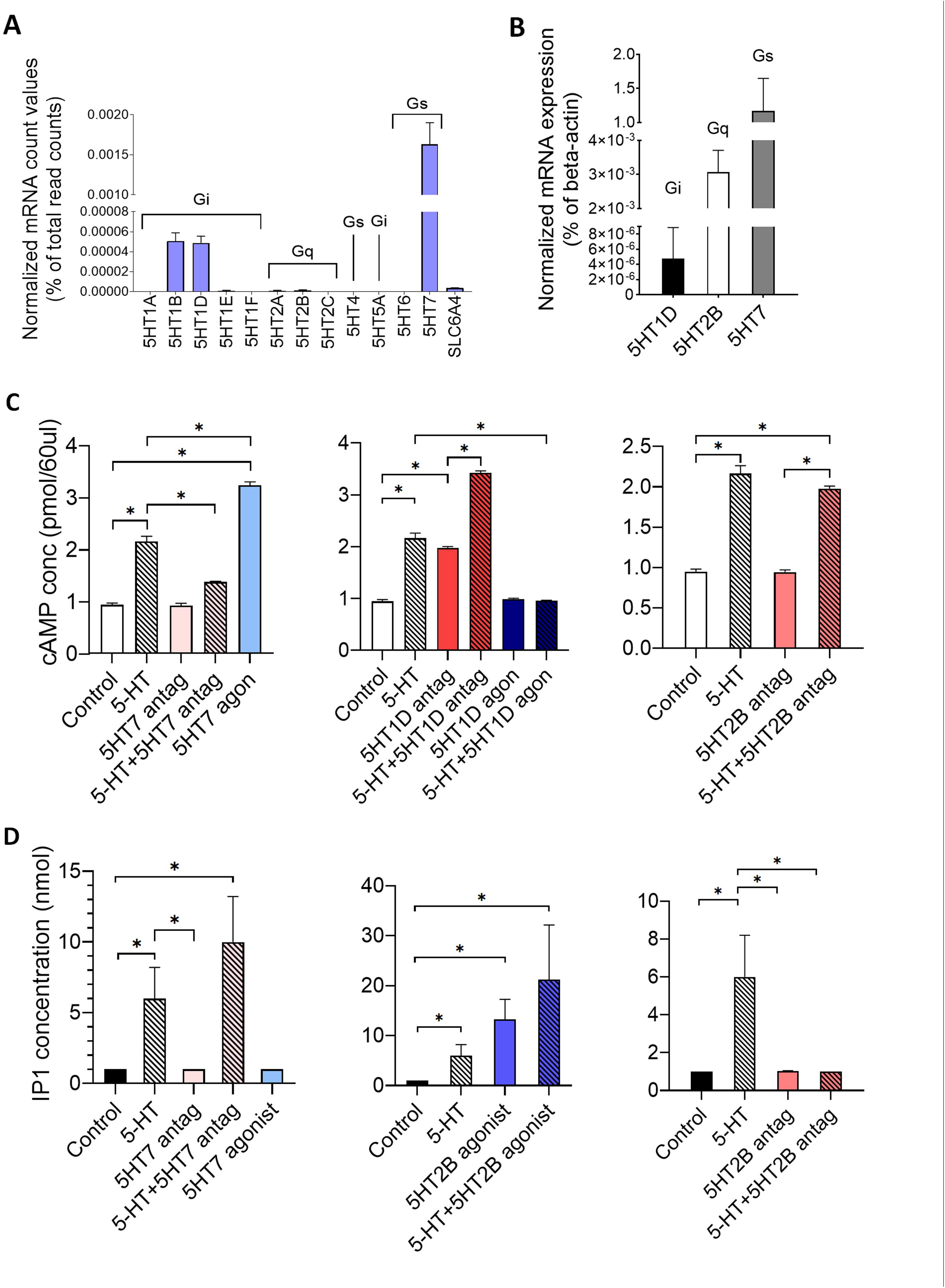
Expression of 5-HT receptors and second messenger response to selective ligands in HME-hTert cells. **A**. Expression pattern of 5-HT receptors in breast epithelial cells based on RNA-Seq data. Read counts of 5-HT receptor isoforms were normalized to total read counts. Associated G-protein are marked for each isoform. **B**. RT-qPCR validation of the expression of selected receptor isoforms for each G-protein coupling (G_i_-HTR1D, G_q_-HTR2B, G_s_-HTR7). mRNA levels were normalized to β-actin. **C**. Intracellular cAMP levels were measured upon 5-HT treatment or selective receptor modulation of the HTR1D, HTR2B and HTR7 isoforms by small molecule agonists (blue bars) and antagonists (red bars). **D**. IP1 levels were quantified as a surrogate for IP3 production following 5-HT or selective receptor agonists (blue bars) and antagonists (red bars) treatments.

Selective activation of 5HTR7 (by LP211), a receptor associated with G_S_, enhanced cAMP production, while the 5HTR7 antagonist SB269970 strongly suppressed cAMP induction by 5-HT (Figure 5C, left panel). 5HTR2B inhibitors (e.g. LY266097) did not reverse the effect of 5-HT (Figure 5C, right panel). In contrast, a 5HTR1D antagonist LY310762 further enhanced 5-HT-induced cAMP levels, while the 5HTR1D agonist tested (PNU142633) blocked the effect of 5-HT (Figure 5C, center graph).

G_q_α-associated receptors activate phospholipase C, generating diacylglycerol and inositol triphosphate (IP3) as second messengers (assessed here by its stable first breakdown product, IP1). The inhibition of 5HTR7 did not decrease 5-HT-induced IP3 levels (Figure 5D, left panel). On the other hand, G_q_α-coupled 5HTR2B receptor activation by the compound BW723C86 stimulated IP3 production, which was further increased by 5-HT (Figure 5D, right panel).

### Anti-proliferative effect of selective 5-HT receptor modulators align with coupled G proteins

To identify receptor isoforms mediating antiproliferative effects, a library of 5-HT receptor modulators was screened (Supplementary figure 4. and Table 1). G_S_-coupled receptor agonists, such as activators of 5HTR7 (IC50 = 3.63 µM) and 5HTR6 (IC50 = 2.99 µM) demonstrated potent antiproliferative effects, while cells did not respond to G_S_-coupled receptor antagonists (Figure 6A,B). Antagonists for G_i_-coupled 5HTR1B (IC50 = 2.97 µM), 5HTR1D (IC50 = 2.54 µM) and 5HTR5A (IC50 = 2.23 µM) markedly reduced cell counts (Figure 6D). On the other hand, agonists for G_i_-coupled receptors (e.g., 5HTR1A, 5HTR1D, 5HTR5A) had no significant impact on proliferation, with the exception of the 5HTR1F agonist (IC50 = 3.249 µM) (Figure 6C). Finally, G_q_-coupled receptor antagonists such as 5HTR2A antagonist (IC50 = 3.71 µM) or LY266097 (5HTR2B antagonist, IC50 = 3.58 µM) markedly reduced cell counts (Figure 6F). From the total of 21 compounds tested, 8 compounds suppressed cell growth at concentrations below 4 μM, including 5 G_i_ or G_q_ inhibitors and 2 G_S_ agonists (Figure 7).

**Table 1.** Selective 5-HT modulator screen of compounds for antiproliferative activity in HME-hTert. Dose-response screening results for a total of 21 compounds tested on HME-hTert cells. Presented data includes target receptors, compound activity, relative inhibitory effect at 10 μM concentration, IC50 values with 95% confidence intervals. IC50 values below 4 μM highlighted in light blue background.

| 5HT-R selective compound | Effect | Receptor coupling | Target receptor | Relative growth at 10µM concentration (% of control) | p-value (10µM) | IC50 value (µM) | 95% confidence interval (µM) |
| --- | --- | --- | --- | --- | --- | --- | --- |
| BIMU8 | Agonist | Gs | 5HT4 | 87,3 | 0.13 | 56.26 | 30.09 - 182.0 |
| WAY181187 |  |  | 5HT6 | 1,2 | <0.001 | 2.99 | 1.40 - 6.47 |
| LP211 |  |  | 5HT7 | 6,6 | <0.001 | 3.63 | 1.60 - 8.38 |
| GR113808 | Antagonist |  | 5HT4 | 87,3 | 0.04 | 49.16 | 30.33 - 101.4 |
| PRX07034 |  |  | 5HT6 | 55,7 | <0.001 | 13.45 | 6.99 - 29.2 |
| SB269970 |  |  | 5HT7 | 85,8 | 0.088 | 46.91 | 24.59 - 152.1 |
| IPSAPIRONE | Agonist | Gi | 5HT1A | 81,8 | 0.077 | 26.71 | 11.13 - 131.0 |
| PNU142633 |  |  | 5HT1D | 92,6 | 0.066 | 31.78 | 21 - 54.23 |
| BRL54443 |  |  | 5HT1E/F | 64,6 | <0.001 | 12.81 | 4.31 - 45.55 |
| LY344864 |  |  | 5HT1F | 1,3 | 0.011 | 3.25 | 1.38 - 7.71 |
| UCSF648 |  |  | 5HT5A | 91 | 0.3 | 46.19 | 27.78 - 99.84 |
| NAD299 | Antagonist |  | 5HT1A | 95,2 | 0.148 | 123.4 | 48.43 - ND |
| SB216641 |  |  | 5HT1B | 0,8 | <0.001 | 2.97 | 1.32 - 6.69 |
| LY310762 |  |  | 5HT1D | 1,7 | <0.001 | 2.54 | 1.25 - 5.16 |
| SB699551 |  |  | 5HT5A | 0,9 | <0.001 | 2.23 | 1.17 - 4.25 |
| BW723C86 | Agonist | Gq | 5HT2B | 74,1 | 0.002 | 27.64 | 14.34 - 75.05 |
| CP809101 |  |  | 5HT2C | 46,7 | 0.001 | 9.014 | 5.58 - 14.88 |
| R96544 | Antagonist |  | 5HT2A | 22,3 | 0.037 | 3.71 | 2.43 - 5.66 |
| LY266097 |  |  | 5HT2B | 17,6 | <0.001 | 3.58 | 2.31 - 5.55 |
| SB204741 |  |  | 5HT2B | 92,9 | 0.345 | 90.02 | 35.08 - ND |
| RS102221 |  |  | 5HT2C | 70,8 | <0.001 | 19.25 | 12.59 - 31.56 |

above 4µM threshold below 4µM threshold

**Figure 6.**
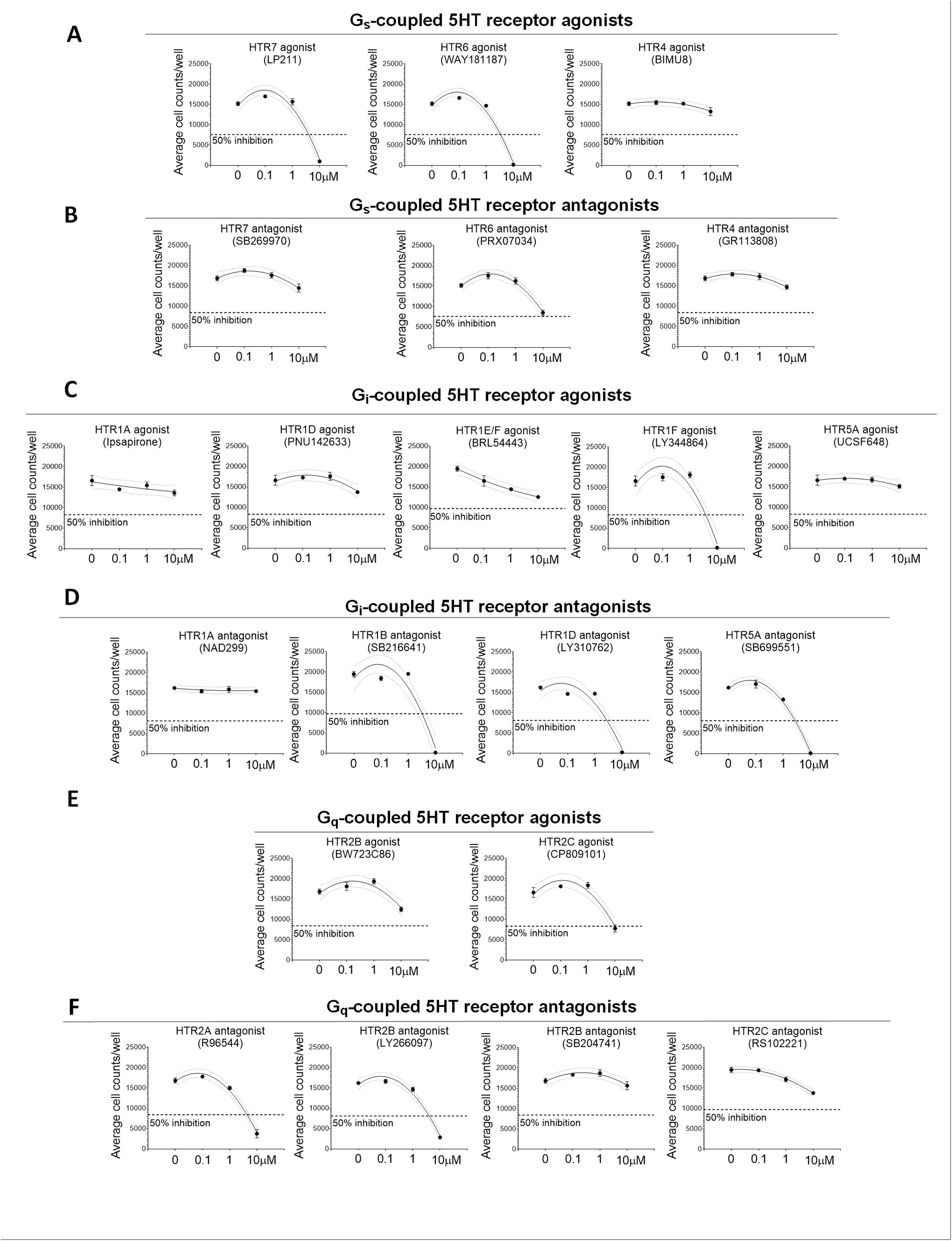
Dose-response curves of the growth suppressive effects of selective 5-HT receptor modulators. Comparison of the antiproliferative effects of selective 5-HT receptor modulators in HME-hTert cells. Dose-response curves represent live cell counts after 96 hours of growth, assessed by microscopy and image analysis. The 21 compounds tested were grouped by their associations with G-proteins. **A**. G_s_-coupled receptor agonists and **B**. antagonists, **C**. G_i_-coupled receptor agonists, **D**. G_i_-coupled receptor antagonists, **E**. G_q_-coupled receptor agonists and **F**. G_q_-coupled receptor antagonists. Standard curves were interpolated for each graph using second-order polynomial (quadratic) interpolation to generate dose-response curves (solid lines) with 95% confidence intervals (dotted lines).

**Figure 7.**
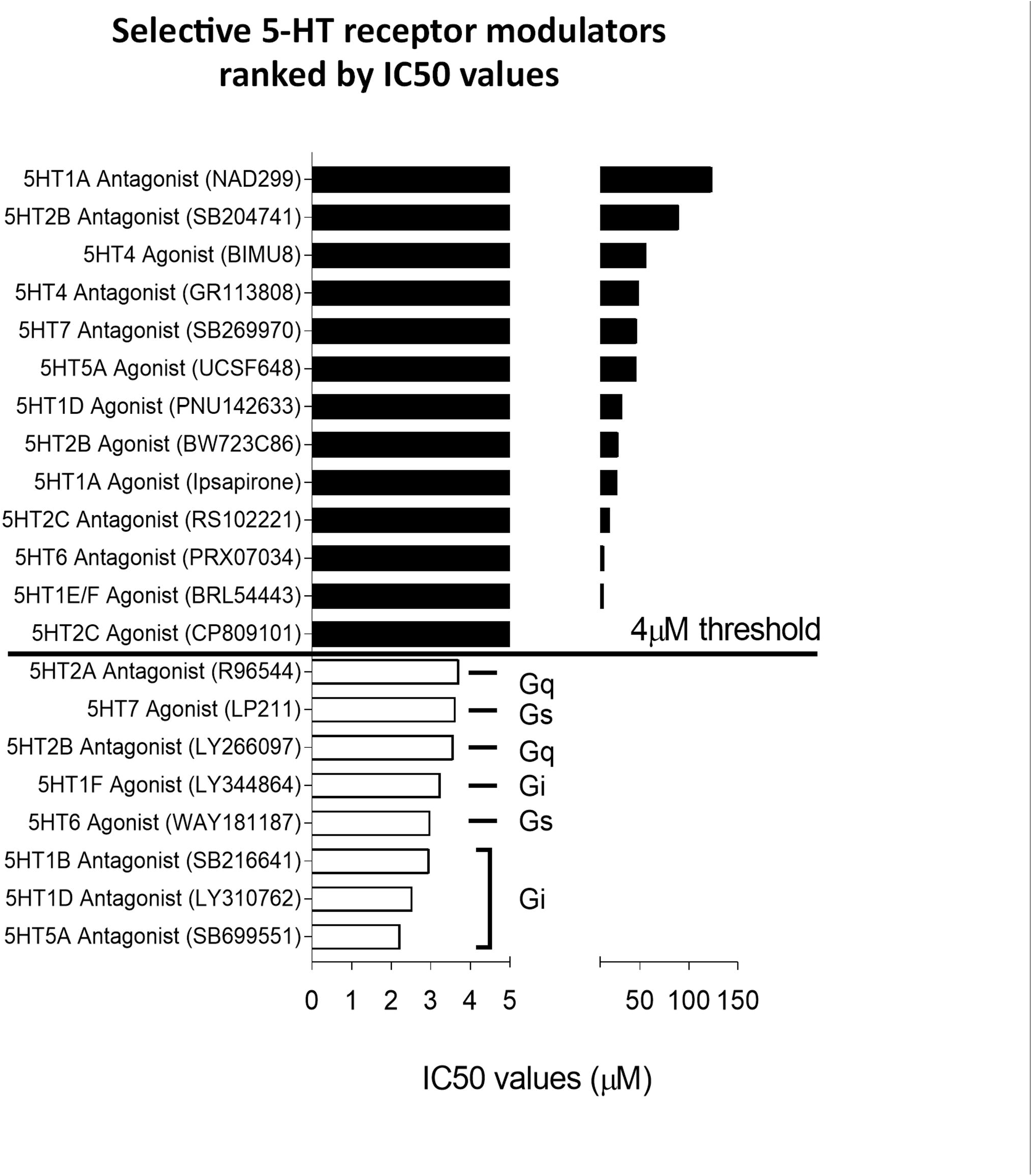
Ranking of selective 5-HT receptor modulators by observed IC50 values for cell growth. Compounds were ranked in decreasing order and labelled based on their G protein coupling. Threshold was defined at 4 µM as a physiologically relevant dose.

To confirm the impact of the 3 most prevalent receptors on breast cell growth, each representing one G protein type, 5-HT or paroxetine effects were modulated by pre-treatment with receptor-selective agonists and antagonists. As previously shown, 5-HT and paroxetine alone or in combination suppressed normal cell proliferation (Figure 8A). Furthermore, antagonists for 5HTR2B (LY266097) and 5HTR1D (LY310762) and the 5HTR7 agonist LP211 showed significant growth suppression (Figure 8B, C, each at 3 µM). In combination treatment the G_S_-coupled 5HTR7 antagonist SB269970, but not the inhibitors for 5HTR2B or 5HTR1D, reversed growth suppression by 5-HT at 10 µM, by paroxetine (1 µM) or 5-HT+paroxetine (Figure 8D). In contrast, the 5HTR7 agonist LP211 enhanced, while the 5HTR1D agonist PNU142633 reversed and the 5HTR2B agonist reduced the antiproliferative effects of paroxetine (Figure 8E). To further emphasize the key role of G_S_-induced cAMP signaling in cell growth, the PKA inhibitor H89 and the adenylyl cyclase inhibitor SQ22536 (both at 3 µM) also blocked the growth suppressive effects of 5-HT and paroxetine (Figure 8F).

**Figure 8.**
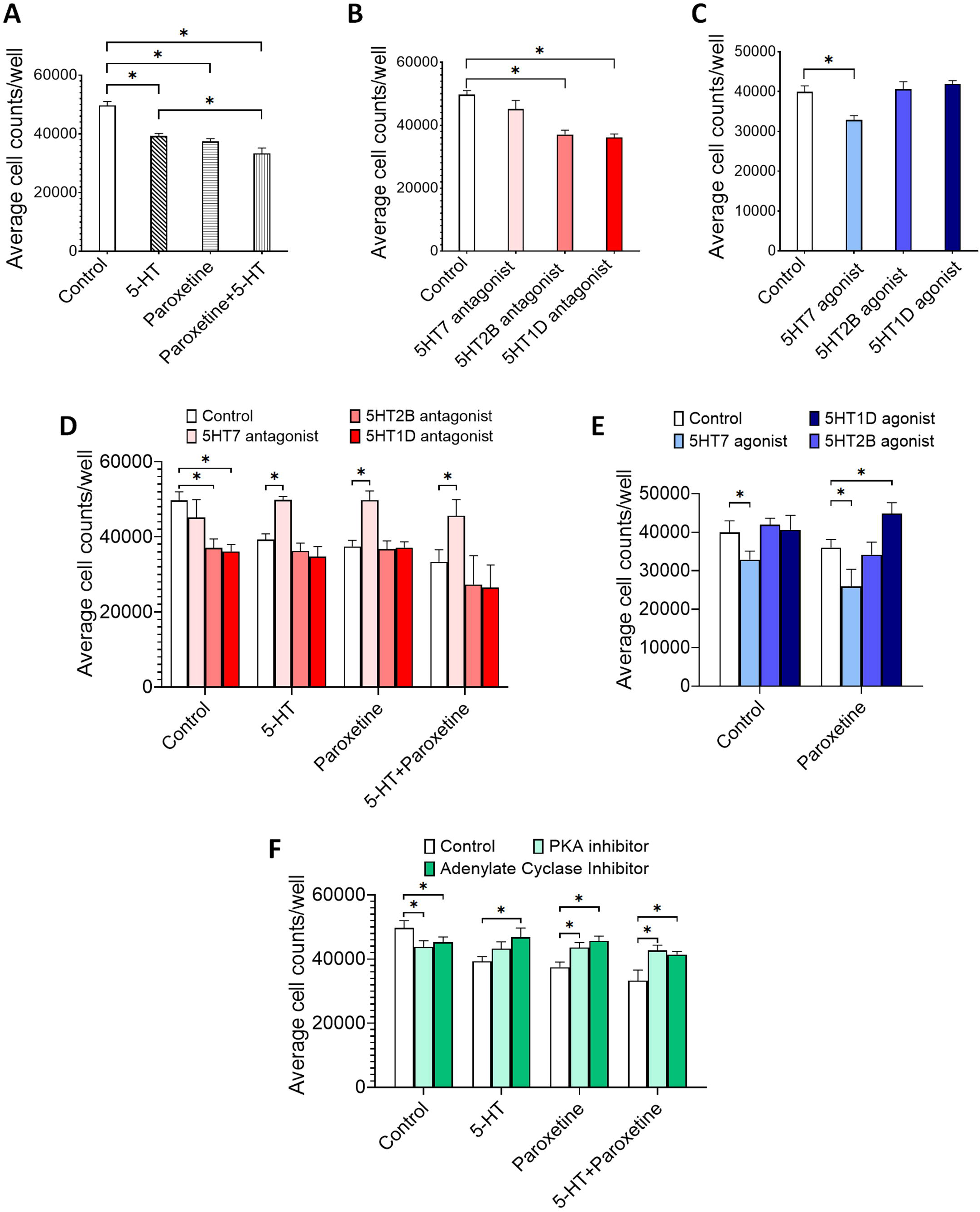
Combining selective 5-HT receptor modulators with 5-HT and/or paroxetine highlights the antiproliferative effect of G_s_-coupled receptor signaling. Comparison of antiproliferative activity of selective 5-HT receptor modulators combined with 5-HT and/or paroxetine in HME-hTert. Cell growth was assessed by 2D microscopy and image analysis-based cell counting upon pre-treatment with either vehicle, 5-HT and/or paroxetine, followed by addition of selective 5-HT receptor modulators. Cell counts shown following a 72 hour treatment by **A**. 5-HT, paroxetine or their combination only, **B**. antagonists selective for 5-HT receptor 7, 2B or 1D, **C**. agonists selective for 5-HT receptor 7, -2B or -1D, **D**. 5-HT, paroxetine or combination after pre-treatment with antagonists of 5HTR7, -2B or -1D, **E**. vehicle or paroxetine after pre-treatment with agonists of 5HTR7, -2B or -1D, **F**. 5-HT, paroxetine or combination following pre-treatment by vehicle, a PKA inhibitor (H89) or an adenylyl cyclase inhibitor (SQ22536).

These findings highlight the G_S_α subunit as a key mediator of serotonergic antiproliferative effects, reversible by G_q_- and G_i_-coupled receptor activation. As 5HTR1D and 5HTR2B represent inhibitory mechanisms to 5-HT-dependent growth suppression, combinations of 5HTR1D and 5HTR2B antagonists with G_S_α activators should be tested for anti-proliferative and tumor suppressive potential.

## Methods

### Cells, Microscopy-Based 2D Proliferation Assay and Compound Screening

Telomerase-immortalized normal breast epithelial cells (HME-hTert) were maintained in MEBM medium supplemented with MEGM growth factor SingleQuots (Lonza, Basel, CH). Cell line authentication was confirmed via deep sequencing prior to experimental use, and regular mycoplasma testing was conducted using a Promokine PCR-based assay. Cell proliferation was assessed using microscopy and image analysis-based assay. HME-hTert cells were seeded at low density in 96-well optical plates and treated for three days. Following treatment, cells were fixed with 4% formaldehyde, and nuclei were labeled with DAPI stain. Imaging was performed using a Leica DMi8 inverted fluorescence microscope, and ImageJ software was used for image analysis with a custom macro incorporating contrast enhancement, nuclear segmentation, and object counting (size range: 5 < X < 400 pixels). Dose-responses to serotonin receptor modulators was screened in the 0.1-10 μM concentration range in 2D proliferation assays. A list of serotonin receptor modulators with chemical structures is shown in supplementary figure 4.

### 3D Epithelial Spheroid Generation and Measurement

HME-hTert cells were embedded in a 1:1 mixture of Matrigel and single-cell suspension in 96-well optical plates. Treatments were initiated on day 6 of spheroid formation and continued for seven days. Phase-contrast images were taken from treated spheroids, and ImageJ analysis was performed to measure spheroid central diameters and spheroid volumes were calculated.

### Immunolabeling, ROS, and γH2AX Measurement

For the quantification of reactive oxygen species (ROS) production after 5-HT or paroxetine treatment cells were stained with the DCFDA reagent at 10 µM final concentration, and counterstained with HOECHST. Cells were pre-treated for 24 hours, followed by DCFDA loading and continued treatment. Microscopic images were taken from live HME-hTert cells 1 hour after DCFDA staining and evaluated using ImageJ, where the area of DCFDA staining was normalized to the number of HOECHST-stained nuclei. Cell fractions positive for double-stranded DNA breaks were quantified by γH2AX immunolabeling. Cells were pre-treated for 1 hour, fixed with 4% formaldehyde, stained for γH2AX and with DAPI. γH2AX positivity relative to the total number of nuclei was calculated and presented for each treatment. Immunostaining was performed as described previously (20).

### Western Blotting

Cell lysates were prepared using RIPA buffer with protease inhibitors and NaVO3 to inhibit phosphatases. Protein quantification was conducted using the Pierce™ BCA protein assay kit (ThermoFisher Scientfic). Proteins were separated by SDS-PAGE, transferred to PVDF membranes, and probed with primary antibodies in 5% BSA-TBST. Washing steps were performed using TBST buffer. Membranes were then probed with secondary antibodies conjugated with DyLight 680 or DyLight 800 fluorophores diluted in 5% BSA-TBST buffer. Detection of signal from membranes was performed using a Licor Odyssey imaging system. Perkin Elmer EnVision 2105 Multi-Mode Microplate Reader

### Measurement of cAMP and IP1 Second Messenger Levels

cAMP levels were measured using the cAMP-Screen Direct™ ELISA (Thermo Fisher, Waltham, MA) assay. Cells treated for 5 minutes were lysed, and lysates were assayed in a 96-well plate with cAMP-AP conjugates and anti-cAMP antibodies. Luminescence was detected using a Synergy LX plate reader (BioTek, Winooski, VA) and concentrations were interpolated from standard curves. IP1 levels were quantified by the HTRF IP-One Gq kit (Revvity Inc, Waltham, MA) using an EnVision 2105 multi-mode microplate reader (Perkin Elmer, Waltham, MA). Treatments were performed in stimulation buffer containing 50mM LiCl_2_. Competitive measurement was conducted by mixing IP1-D2 and anti-IP1 cryptate reagents with cell lysates. Fluorescent signals were detected at 665nm and 615nm, based on which HTRF ratios were calculated.

### RT-qPCR Measurements

Transcripts for the 5HTR1D, 5HTR2B and 5HTR7 serotonin receptor isoforms were transcribed using gene-specific reverse primers and reverse transcriptase, and cDNAs were amplified in qPCR assays with TaqMan probes. mRNA copy numbers were interpolated from standard curves derived from the measurement of serial dilution series of single-stranded, transcript-specific amplicon standards. Expression levels of the selected 5-HT receptor isoforms were normalized to β-actin expression and presented as percentages.

### Bulk RNA Sequencing and Gene Set Enrichment Analysis (GSEA)

RNA was extracted from treated HME-hTert cells using Trizol reagent. Quality was assessed using the Agilent Bioanalyzer. RNA sequencing was performed on a DNBSEQ-G400 platform. Sequencing data was processed on the Galaxy platform, including trimming (Trimmomatic), alignment (HISAT2), quantification (FeatureCounts).

GSEA was conducted using the GSEA software versionv4.3.3 [build: 16] to identify enriched gene sets in the transcriptomic dataset comparing vehicle-treated and paroxetine-treated samples. Raw RNA sequencing files are uploaded to NCBI and accessible by the accession number PRJNA1210992 (https://www.ncbi.nlm.nih.gov/bioproject/PRJNA1210992/). The analysis utilized the c2.cp.reactome-v2024.1.Hs.symbols.gmt gene set database from MSigDB, with 1000 gene set permutations for statistical significance. The expression dataset contained 25702 genes across six samples, annotated with the Human NCBI gene ID MSigDB v2024.1 chip platform. Phenotypic labels defined 2 groups and genes were ranked using the Signal2Noise metric, sorted in descending order. Collapsing mode was set to “Max_probe”, and gene set sizes were restricted to 15-500 genes. Normalization was performed by selecting “meandiv”, alongside with “timestamp” selection as the permutation seed and “no_balance” selection for randomization. Reactome pathways identified from the comparison of vehicle-or paroxetine-treated cells are presented in figures 3-4.

### Statistical Evaluations

Data are expressed as mean ± SEM. Experiments included at least three independent biological replicates. Statistical comparisons between two groups were performed using unpaired Student’s t-test. ANOVA with Mann-Whitney post-hoc test was used for multiple group comparisons. A p-value < 0.05 was considered statistically significant.

## Discussion

Emerging evidence demonstrate altered functionality of monamine neurotransmitter signaling systems on multiple levels in the course of malignant transformation (8, 21). In the case of serotonin, carcinogenesis changes its biosynthesis and tissue levels, 5-HT receptor expression profiles, and induces a shift to glycolytic metabolism (11, 14, 22). By inhibiting the uptake of serotonin in cells expressing the SERT transporter, SSRIs modulate 5-HT levels in the extracellular space, and thus the exposure of 5HTRs to the neurotransmitter. Epidemiologic data from the TCGA indicate that 3-4% of breast tumors may leverage their progression by upregulating SERT through gene amplification. High expression of SERT is associated with a deterioration in overall survival, which is in agreement with elevated 5-HT levels proposed as predictive biomarkers in breast cancer (11). In this study we found that SSRIs moderately reduce the proliferation rates of HME-hTert cells. As antidepressants are frequently used drugs in cancer patients, safety has been in the forefront of interest. Clinical assessment of their potential impact on tumor formation and outcome revealed little evidence of increased breast cancer risk associated with the use of SSRIs (23, 24), and there was no correlation with breast cancer mortality (25). However, there was a 36% reduction in breast cancer risk among paroxetine hydrochloride users (26). As paroxetine was reported to show the highest affinity to SERT among SSRIs, the growth suppressive activity of paroxetine surpassing that of other antidepressants tested was not surprizing, and justified its use in the study (27). Given the enhanced anti-proliferative potential of 5-HT by paroxetine in non-malignant mammary epithelial cells, further studies evaluating the use of paroxetine in *in vivo* models of primary breast cancer prevention is warranted.

Several serotonin receptors were shown to contribute to proliferative signaling and cell cycle regulation in breast cancer cells, and in some cases identified as instrumental in the progression of cancer cells (3, 28-30). In MCF-7 cells 5HTR2A was shown to transmit the mitogenic effect of serotonin (31). 5HTR1D was demonstrated to be key to breast cancer cell growth. The knock-down of 5HTR1D suppressed tumor cell proliferation (32). In TNBC cell lines 5HTR7 was upregulated and suppression of 5HTR7 reduced eEF2 kinase and cyclin D1 expression, and inhibited TNBC cell proliferation (33). Nevertheless, considering the co-expression of several 5-HT receptors, the changes evoked by 5-HT represent a compound effect resulting from the activation of all 5HTRs expressed in the cell in its actual state, thus understanding the relative contribution of each receptor has been a challenging task.

In non-transformed breast tissue 5HTR activation leads to markedly different outcomes, including negative regulation of milk synthesis and secretion, involution, and stimulation of a calcium-mobilizing hormone (6). However, the receptor-specific impact of 5-HT on growth regulation in non-cancerous mammary epithelial cells is less elucidated and therefore became the focus of this work. One of the cornerstones of primary cancer prevention in the mammary glands is the suppression of proliferation rates in the epithelial cell layer, to reduce the odds of malignant transformation and carcinogenesis (34). We show that in contrast to breast cancer cell response, 5-HT is inhibitory to normal cell growth (3). In HME-hTert cells 5-HTR7 agonists induced growth arrest. Based on gene expression data and functional responses through second messengers, normal breast cells express receptors covering all 3 major classess of G proteins, hence it was not clear what determines the antiproliferative response to 5-HT. Selective modulation of these pathways by targeting individual receptors allowed us to achieve far greater antiproliferative action than by the universal ligand 5-HT. Therefore, elucidating which receptors activate and which ones block inhibitory signals for growth may highlight novel avenues for more efficient cancer preventive strategies.

Interestingly, none of the 5-HT receptor agonists or inhibitors stimulated proliferation on their own, but reversed or enhanced 5-HT-dependent growth suppression. We therefrom conclude that in non-malignant cells 5HTR activation only serves as a controlling, limiting factor, but not a growth signal itself.

All 5HTRs interact with one of 3 separate heterotrimeric G proteins, utilizing two distinct downstream signaling pathways (35). Our systematic comparison of the anti-proliferative effects of receptor selective agonists and antagonists suggest that it is the stimulation of the G_S_α and the suppression of G_i_ or G_q_ –dependent signaling, which can amount to radical growth reduction in non-malignant mammary epithelial cells. A relatively high frequency of gene amplifications occurring in breast tumors may indicate addiction of breast cancer cells to G_S_α (Supplementary figure 5). Conversely, alterations in G_i_ or G_q_α, 5HTR 1D, 1F, 2B or 7 are rare, possibly due to the expression of a diverse array of 5HTRs in most cells making a clearly pro-tumorigenic signaling less likely. Therefore, heterotrimeric Gα subunits might be suitable drug targets in such cases where a high level of GPCR redundancy complicates effective receptor-specific blockade (36). As G-protein isoform-specific signaling appears to dictate proliferation response to serotonergic analogues, the potential synergy between G_S_α agonists and G_q_α or G_i_α antagonists may be a promising target for further investigations.

The limitations of the study include the potential for many ligands to cross-react with related receptor subtypes which may present with off-target effects. G_q_-coupled receptor ligands, particularly some selective for 5HTR2, exhibit biased agonism through beta-catenin activation (37). Furthermore, our study cannot take into account receptor-independent effects of the changes in serotonin levels. Apart from its receptor-mediated actions, 5-HT is a substrate for posttranslational modifications of select proteins. Serotonylation was shown to promote thrombocyte activation or influence epigenetic programming through histone serotonylation and driving ependymoma tumorigenesis (38-40). These exciting aspects of 5-HT biology will demand further attention.

In summary, complementing previous research, our study raises awareness to the therapeutic opportunities associated with selective modulation of 5-HT receptor pathways (14, 16). Repurposing pharmaceuticals modulating neurotransmitter pathways remains a promising way to improve cancer therapy and enable chemoprevention (41).

## Acknowledgements

The authors thank Dr. Attila Szőllősi for his kind help with the quantitation of IP1 levels using a 5HTRF reader and Dr. Szilárd Póliska for his technical support with next generation sequencing. These studies were funded by the grant K129218 from the National Research, Development and Innovation Office of Hungary to IPU. The research was kindly supported by the University of Debrecen Scientific Research Bridging Fund (DETKA), the UD Publication Support Program and the UD Clinical Center. Also supported by the ÚNKP-21-3 New National Excellence program of the Ministry for Innovation and Technology from the source of the National Research, Development and Innovation Fund to ML.

## Authorship

Contribution: I.P.U. designed the study, conducted some experiments and interpreted the results. M.L. performed most of the experiments, analyzed results. Writing the manuscript – M.L., I.P.U. Review and Editing, I.P.U., P.Á. and M.L. Funding acquisition by I.P.U and P.Á.

## References

1. Jones LA, Sun EW, Martin AM, Keating DJ. The ever-changing roles of serotonin. Int J Biochem Cell Biol. 2020;125:105776.

2. Ray RS, Corcoran AE, Brust RD, Kim JC, Richerson GB, Nattie E, et al. Impaired respiratory and body temperature control upon acute serotonergic neuron inhibition. Science. 2011;333(6042):637–42.

3. Pai VP, Marshall AM, Hernandez LL, Buckley AR, Horseman ND. Altered serotonin physiology in human breast cancers favors paradoxical growth and cell survival. Breast Cancer Res. 2009;11(6):R81.

4. Gautam J, Banskota S, Regmi SC, Ahn S, Jeon YH, Jeong H, et al. Tryptophan hydroxylase 1 and 5-HT7 receptor preferentially expressed in triple-negative breast cancer promote cancer progression through autocrine serotonin signaling. Mol Cancer. 2016;15(1):75.

5. Stapel B, Melzer C, von der Ohe J, Hillemanns P, Bleich S, Kahl KG, et al. Effect of SSRI exposure on the proliferation rate and glucose uptake in breast and ovary cancer cell lines. Sci Rep. 2021;11(1):1250.

6. Marshall AM, Hernandez LL, Horseman ND. Serotonin and serotonin transport in the regulation of lactation. J Mammary Gland Biol Neoplasia. 2014;19(1):139–46.

7. Horseman ND, Collier RJ. Serotonin: a local regulator in the mammary gland epithelium. Annu Rev Anim Biosci. 2014;2:353–74.

8. Kannen V, Bader M, Sakita JY, Uyemura SA, Squire JA. The Dual Role of Serotonin in Colorectal Cancer. Trends Endocrinol Metab. 2020;31(8):611–25.

9. Soto-Pantoja DR, Gaber M, Arnone AA, Bronson SM, Cruz-Diaz N, Wilson AS, et al. Diet Alters Entero-Mammary Signaling to Regulate the Breast Microbiome and Tumorigenesis. Cancer Res. 2021;81(14):3890–904.

10. Frobe A, Cicin-Sain L, Jones G, Soldic Z, Lukac J, Bolanca A, et al. Plasma free serotonin as a marker for early detection of breast cancer recurrence. Anticancer Res. 2014;34(3):1167–9.

11. Xie QE, D. X, Wang M, Xie F, Zhang Z, Cao Y, et al. Identification of Serotonin as a Predictive Marker for Breast Cancer Patients. Int J Gen Med. 2021;14:1939–48.

12. Peng Y, Gu J, Liu F, Wang P, Wang X, Si C, et al. Integrated analysis of microbiota and gut microbial metabolites in blood for breast cancer. mSystems. 2024;9(11):e0064324.

13. McCorvy JD, Roth BL. Structure and function of serotonin G protein-coupled receptors. Pharmacol Ther. 2015;150:129–42.

14. Ballou Y, Rivas A, Belmont A, Patel L, Amaya CN, Lipson S, et al. 5-HT serotonin receptors modulate mitogenic signaling and impact tumor cell viability. Mol Clin Oncol. 2018;9(3):243–54.

15. Jose J, Tavares CDJ, Ebelt ND, Lodi A, Edupuganti R, Xie X, et al. Serotonin Analogues as Inhibitors of Breast Cancer Cell Growth. ACS Med Chem Lett. 2017;8(10):1072–6.

16. Warchal SJ, Dawson JC, Shepherd E, Munro AF, Hughes RE, Makda A, et al. High content phenotypic screening identifies serotonin receptor modulators with selective activity upon breast cancer cell cycle and cytokine signaling pathways. Bioorg Med Chem. 2020;28(1):115209.

17. Ashbury JE, Levesque LE, Beck PA, Aronson KJ. Selective Serotonin Reuptake Inhibitor (SSRI) Antidepressants, Prolactin and Breast Cancer. Front Oncol. 2012;2:177.

18. Balakrishna P, George S, Hatoum H, Mukherjee S. Serotonin Pathway in Cancer. Int J Mol Sci. 2021;22(3).

19. Moller IR, Slivacka M, Nielsen AK, Rasmussen SGF, Gether U, Loland CJ, et al. Conformational dynamics of the human serotonin transporter during substrate and drug binding. Nat Commun. 2019;10(1):1687.

20. Lengyel M, Molnar A, Nagy T, Jdeed S, Garai I, Horvath Z, et al. Zymogen granule protein 16B (ZG16B) is a druggable epigenetic target to modulate the mammary extracellular matrix. Cancer Sci. 2025;116(1):81–94.

21. Jayachandran P, Battaglin F, Strelez C, Lenz A, Algaze S, Soni S, et al. Breast cancer and neurotransmitters: emerging insights on mechanisms and therapeutic directions. Oncogene. 2023;42(9):627–37.

22. Sola-Penna M, Paixao LP, Branco JR, Ochioni AC, Albanese JM, Mundim DM, et al. Serotonin activates glycolysis and mitochondria biogenesis in human breast cancer cells through activation of the Jak1/STAT3/ERK1/2 and adenylate cyclase/PKA, respectively. Br J Cancer. 2020;122(2):194–208.

23. Coogan PF, Strom BL, Rosenberg L. SSRI use and breast cancer risk by hormone receptor status. Breast Cancer Res Treat. 2008;109(3):527–31.

24. Eom CS, Park SM, Cho KH. Use of antidepressants and the risk of breast cancer: a meta-analysis. Breast Cancer Res Treat. 2012;136(3):635–45.

25. Wernli KJ, Hampton JM, Trentham-Dietz A, Newcomb PA. Use of antidepressants and NSAIDs in relation to mortality in long-term breast cancer survivors. Pharmacoepidemiol Drug Saf. 2011;20(2):131–7.

26. Wernli KJ, Hampton JM, Trentham-Dietz A, Newcomb PA. Antidepressant medication use and breast cancer risk. Pharmacoepidemiol Drug Saf. 2009;18(4):284–90.

27. Hyttel J. Comparative pharmacology of selective serotonin re-uptake inhibitors (SSRIs). Nordic Journal of Psychiatry. 1993;47(Sup30):5-12.

28. Marin P, Bécamel C, Chaumont-Dubel S, Vandermoere F, Bockaert J, Claeysen S. Classification and signaling characteristics of 5-HT receptors: toward the concept of 5-HT receptosomes. Hbk Behav Neurosci. 2020;31:91–120.

29. Bockaert J, Claeysen S, Dumuis A, Marin P. Classification and Signaling Characteristics of 5-HT Receptors. Hbk Behav Neurosci. 2010;21:103–21.

30. Ambrosio MR, Magli E, Caliendo G, Sparaco R, Massarelli P, D’Esposito V, et al. Serotoninergic receptor ligands improve Tamoxifen effectiveness on breast cancer cells. BMC Cancer. 2022;22(1):171.

31. Sonier B, Arseneault M, Lavigne C, Ouellette RJ, Vaillancourt C. The 5-HT2A serotoninergic receptor is expressed in the MCF-7 human breast cancer cell line and reveals a mitogenic effect of serotonin. Biochem Biophys Res Commun. 2006;343(4):1053–9.

32. Liu S, He M, Sun H, Wu Y, Jin W. 5-Hydroxytryptamine G-Protein-Coupled Receptor Family Genes: Key Players in Cancer Prognosis, Immune Regulation, and Therapeutic Response. Genes (Basel). 2024;15(12).

33. Cinar V, Hamurcu Z, Guler A, Nurdinov N, Ozpolat B. Serotonin 5-HT7 receptor is a biomarker poor prognostic factor and induces proliferation of triple-negative breast cancer cells through FOXM1. Breast Cancer. 2022;29(6):1106–20.

34. Dunn BK, Kramer BS. Cancer Prevention: Lessons Learned and Future Directions. Trends Cancer. 2016;2(12):713–22.

35. Oldham WM, Hamm HE. Heterotrimeric G protein activation by G-protein-coupled receptors. Nat Rev Mol Cell Biol. 2008;9(1):60–71.

36. Kostenis E, Pfeil EM, Annala S. Heterotrimeric G(q) proteins as therapeutic targets? J Biol Chem. 2020;295(16):5206–15.

37. McCorvy JD, Wacker D, Wang S, Agegnehu B, Liu J, Lansu K, et al. Structural determinants of 5-HT(2B) receptor activation and biased agonism. Nat Struct Mol Biol. 2018;25(9):787–96.

38. Bader M. Serotonylation: Serotonin Signaling and Epigenetics. Front Mol Neurosci. 2019;12:288.

39. Zhao S, Chuh KN, Zhang B, Dul BE, Thompson RE, Farrelly LA, et al. Histone H3Q5 serotonylation stabilizes H3K4 methylation and potentiates its readout. Proc Natl Acad Sci U S A. 2021;118(6).

40. Chen HC, He P, McDonald M, Williamson MR, Varadharajan S, Lozzi B, et al. Histone serotonylation regulates ependymoma tumorigenesis. Nature. 2024;632(8026):903–10.

41. Fu L, Jin W, Zhang J, Zhu L, Lu J, Zhen Y, et al. Repurposing non-oncology small-molecule drugs to improve cancer therapy: Current situation and future directions. Acta Pharm Sin B. 2022;12(2):532–57.

