## Supplementary figures and images for "G protein subtype preference dictates paroxetine-enhanced serotonin receptor response in normal breast epithelial cells"

### Supplemental Figure 1

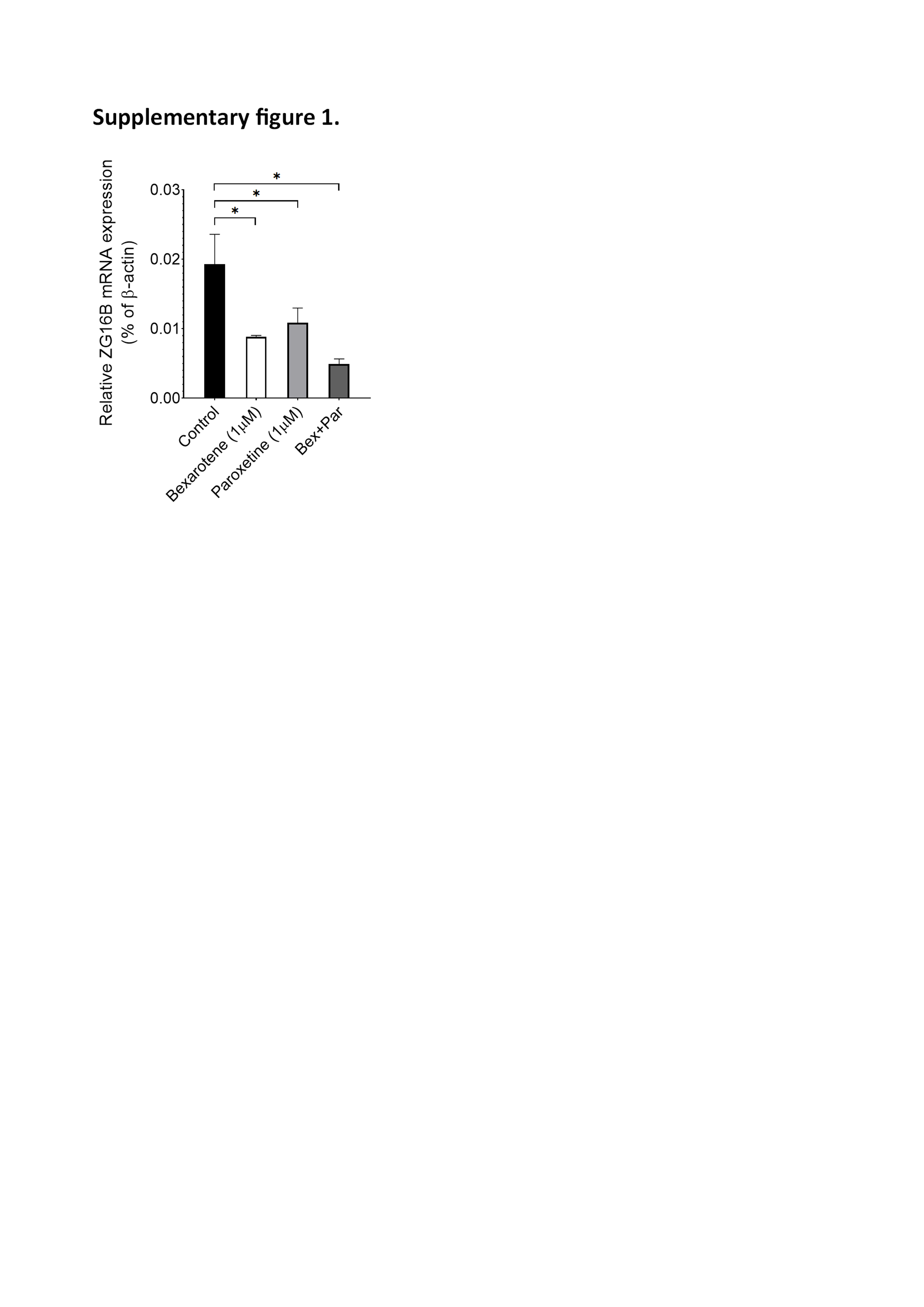

### Supplemental Figure 2

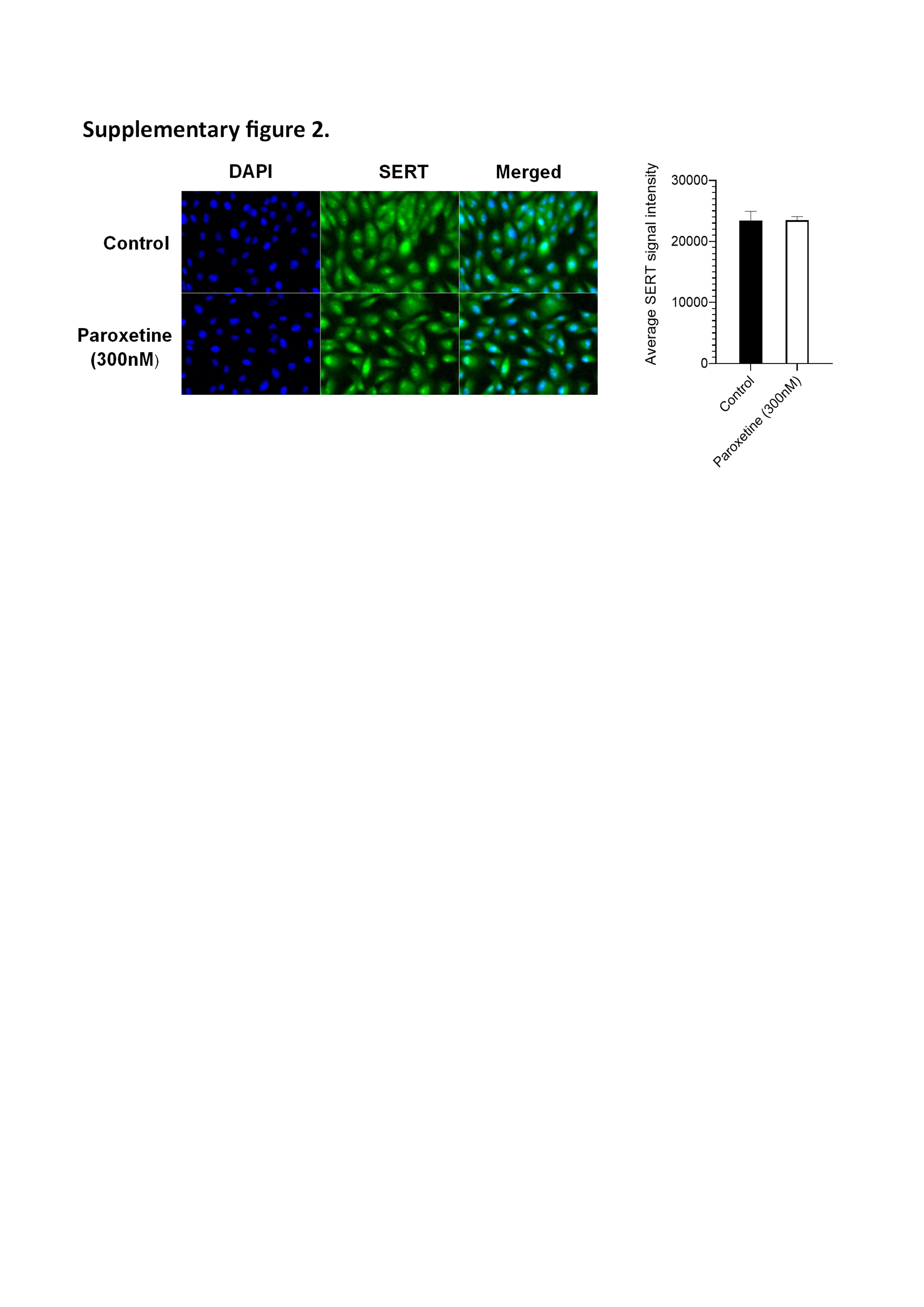

### Supplemental Figure 3

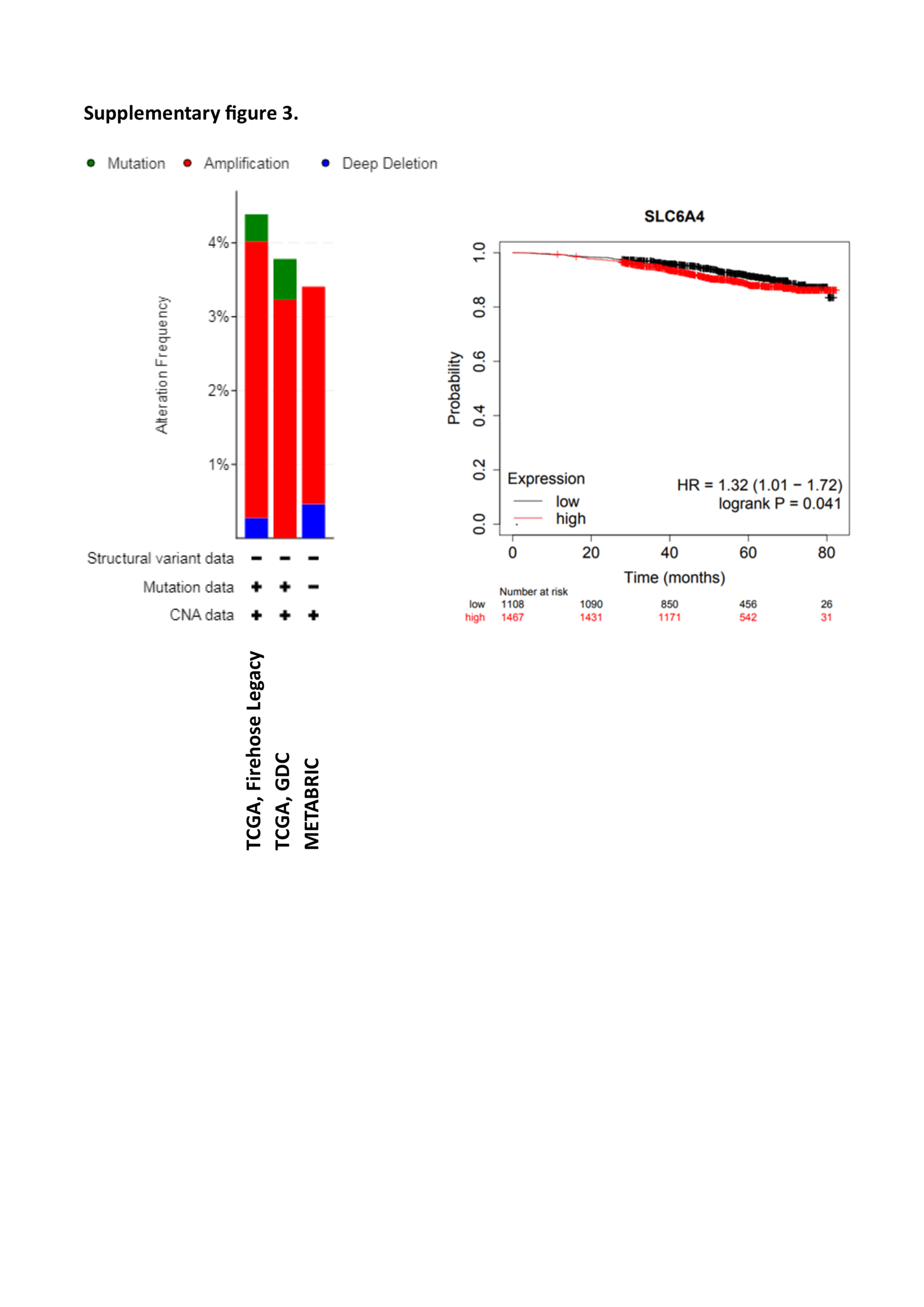

### Supplemental Figure 4

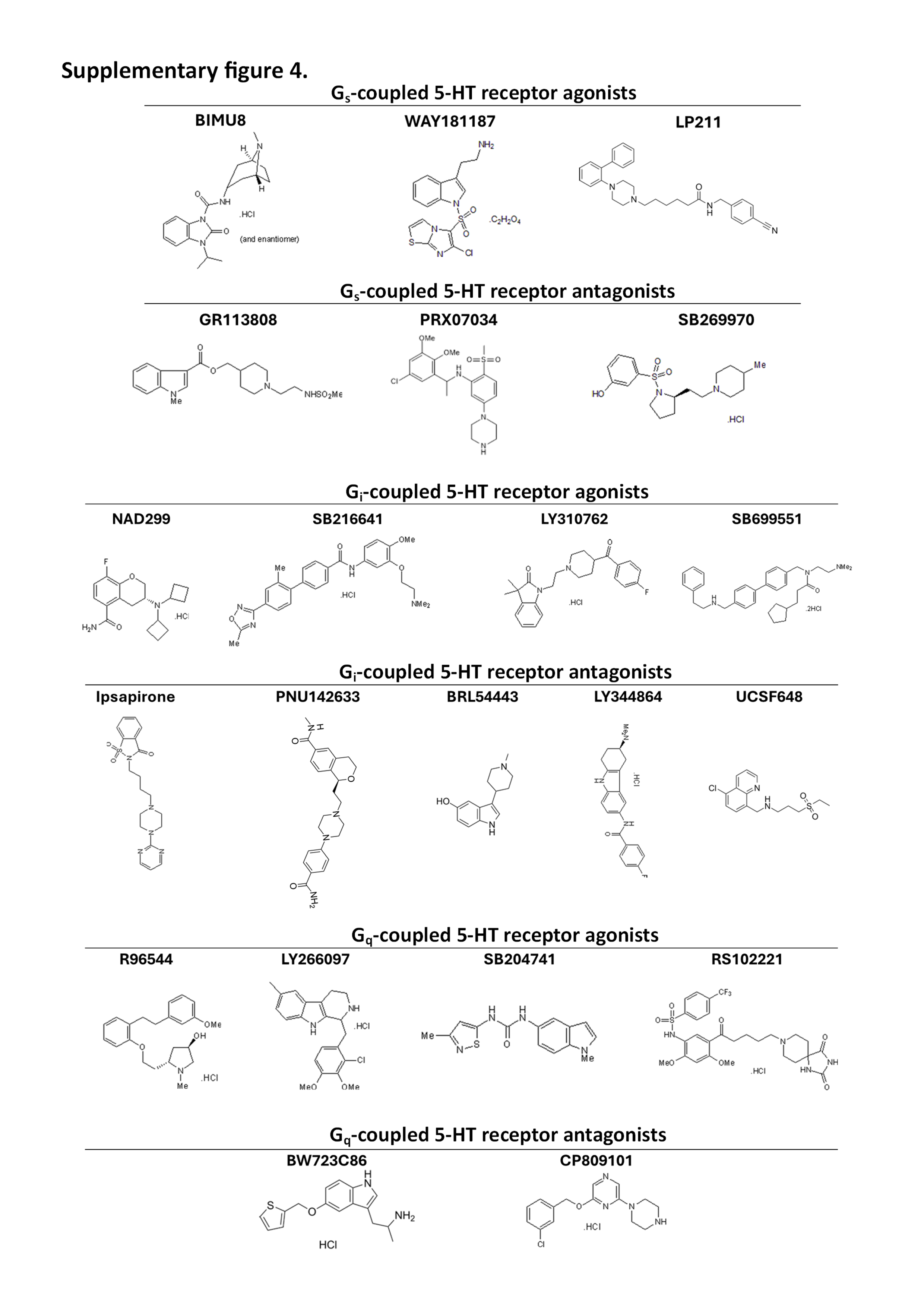

### Supplemental Figure 5

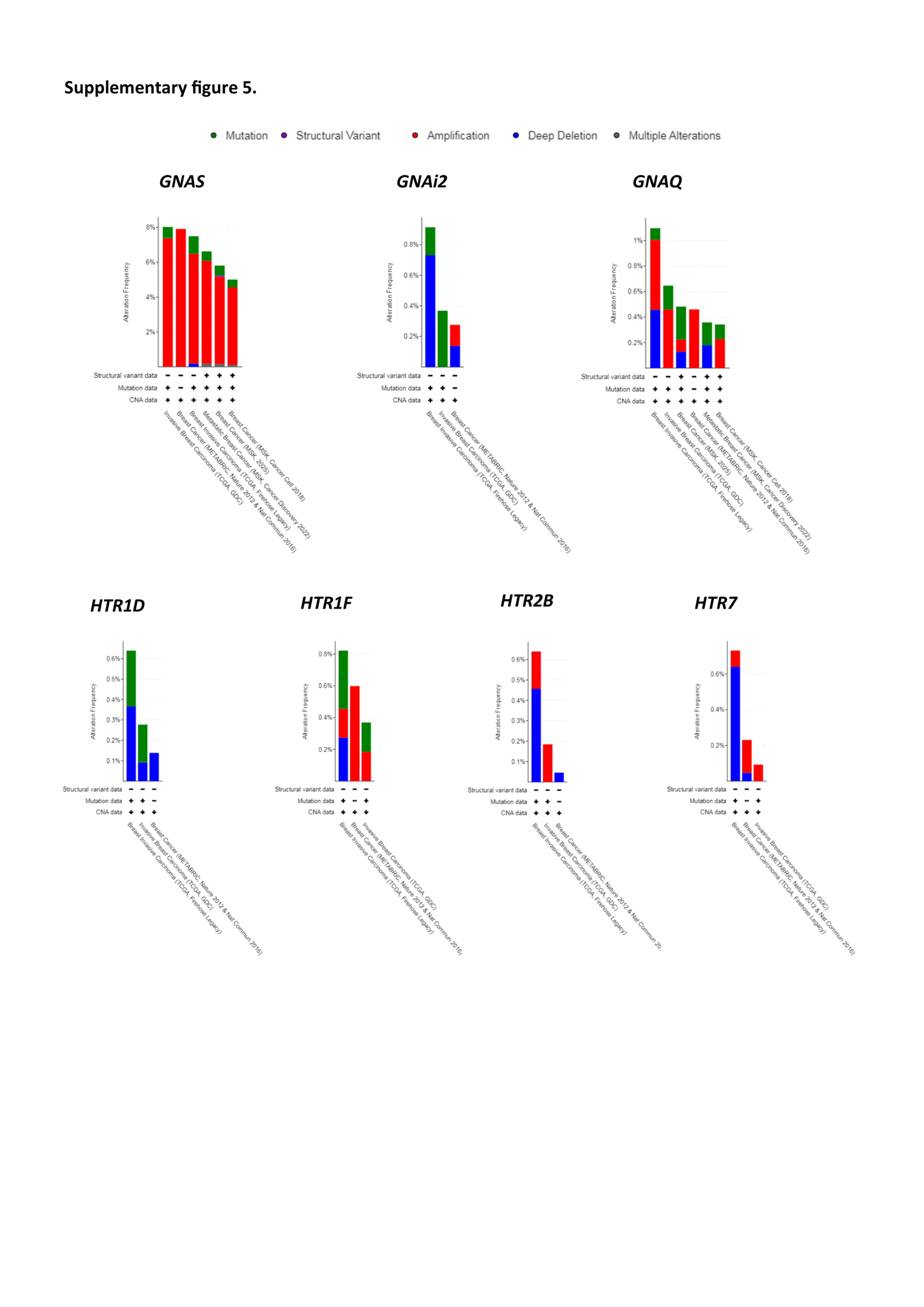
